# Identifying the essential genes of *Mycobacterium avium* subsp. *hominissuis* with Tn-Seq using a rank-based filter procedure

**DOI:** 10.1101/708495

**Authors:** William M. Matern, Robert L. Jenquin, Joel S. Bader, Petros C. Karakousis

## Abstract

*Mycobacterium avium* (Mav) is increasingly recognized as a significant cause of morbidity, particularly in elderly patients or those with immune deficiency or underlying structural lung disease. Generally, Mav infection is treated with 2-3 antimicrobial drugs for at least 12 months. Identification of genes essential for Mav growth may yield novel strategies for improving curative therapy. We have generated saturating genome-wide transposon mutant pools in a commonly used laboratory strain of *Mycobacterium avium* subsp. *hominissuis* (MAC109) and developed a computational technique for classifying annotated genomic features as essential (ES), growth defect (GD), growth advantage (GA), or no-effect (NE) based on the *in vitro* effect of disruption by transposon. We identified 270 features as ES with 230 of these overlapping with ES features in *Mycobacterium tuberculosis*. These results may be useful for identifying drug targets or for informing studies requiring genetic manipulation of *Mycobacterium avium*, which should seek to avoid disrupting ES features to ensure bacterial viability.

**Importance:** *Mycobacterium avium subsp. hominissuis* is an emerging cause of morbidity in vulnerable populations in many countries. It is known to be particularly difficult to treat, often requiring years of antibiotic therapy. In this study we report the genes of *Mycobacterium avium* subsp. *hominissuis* that are required for the organism to grow *in vitro*. Our findings may help guide future research into identifying new drugs to improve the treatment of this serious infection.

## Introduction

The genus *Mycobacteria* contains a variety of difficult-to-treat pathogens, frequently associated with pulmonary disease. One of these pathogens, *Mycobacterium avium* (Mav), is an opportunistic pathogen associated with significant morbidity in the elderly and in patients with underlying lung disease^1,2^ as well as increased mortality in patients with AIDS^3^. Similar to other mycobacteria, Mav is often difficult to treat effectively with existing antibiotic combinations. Current antibiotic regimens require a median of 5 months to convert the sputum to a culture-negative state^4^, with current guidelines suggesting these infections be treated for at least 1 year after sputum conversion^5^. Furthermore, a large fraction of patients fail to convert after 1 year of therapy^4^. Patients could greatly benefit from new therapeutic approaches with greater efficacy and reduced duration.

Transposon sequencing (e.g., TraDIS^6^, Tn-Seq^7^, INseq^8^) has been extensively used to profile haploid genomes and identify gene disruptions that affect bacterial growth under various conditions. Of potential interest in drug development are those drug targets which profoundly disrupt growth on rich media (i.e., “essential” genes). In the current study, we have identified genes critical for Mav growth *in vitro* with the goal of informing future research in Mav pathogenesis and drug development. In order to make gene essentiality predictions, we developed a new statistical approach for calling genes based on ranking the read counts from each mutant and applied this to new Tn-Seq data. We report our predictions of the essential genes of Mav and compare these with the predicted set of essential genes in the closely related human pathogen, *Mycobacterium tuberculosis*.

## Results

### Constructing Genome-wide Transposon Mutant Pools in *Mycobacterium avium*

To identify a suitable strain of Mav for genome-wide mutagenesis, we evaluated the ability of the Himar1 transposon (delivered via ΦmycomarT7^9^), which inserts randomly into thymine-adenine dinucleotide (TA sites), to transform common laboratory strains. Transformation efficiency and spontaneous resistance rate (background) were estimated via CFU counts and are provided in Table S1. Of the 5 strains tested, MAC109 was observed to have the highest transformation efficiency with only ~1% background. Therefore, we decided to proceed with transposon mutagenesis with this strain. Upon transformation, we estimated each of our five independent MAC109 transposon mutant libraries contained between 2.2 – 4.4 × 10^5^ unique insertion events, for a combined total of 1.2 × 10^6^ unique events with ~2% background. To assist with analysis, we recently provided the genome of this strain which was found to contain a 5,188,883 bp chromosome and two plasmids (pMAC109a and pMAC109b) of lengths 147,100 bp and 16,516 bp, respectively^10^. There were 60,129 unique TA sites across the entire genome.

### Confirmation of site bias

It was previously shown that the Himar1 transposon/transposase system has a reduced rate of insertion in sites containing the sequence motif [CG]GNTANC[CG]^7^. Indeed, our results confirm that insertion into these low permissibility sites is much less likely than other sites (Figure 1). Although our approach was able to disrupt nearly all possible insertion sites in the genome not matching this motif (i.e., achieving saturation), a substantial fraction of the low permissibility sites in the chromosome were unoccupied in all five libraries. This effect was less apparent in the plasmids, likely due to their multiple copy number^10^.

**Figure 1:**
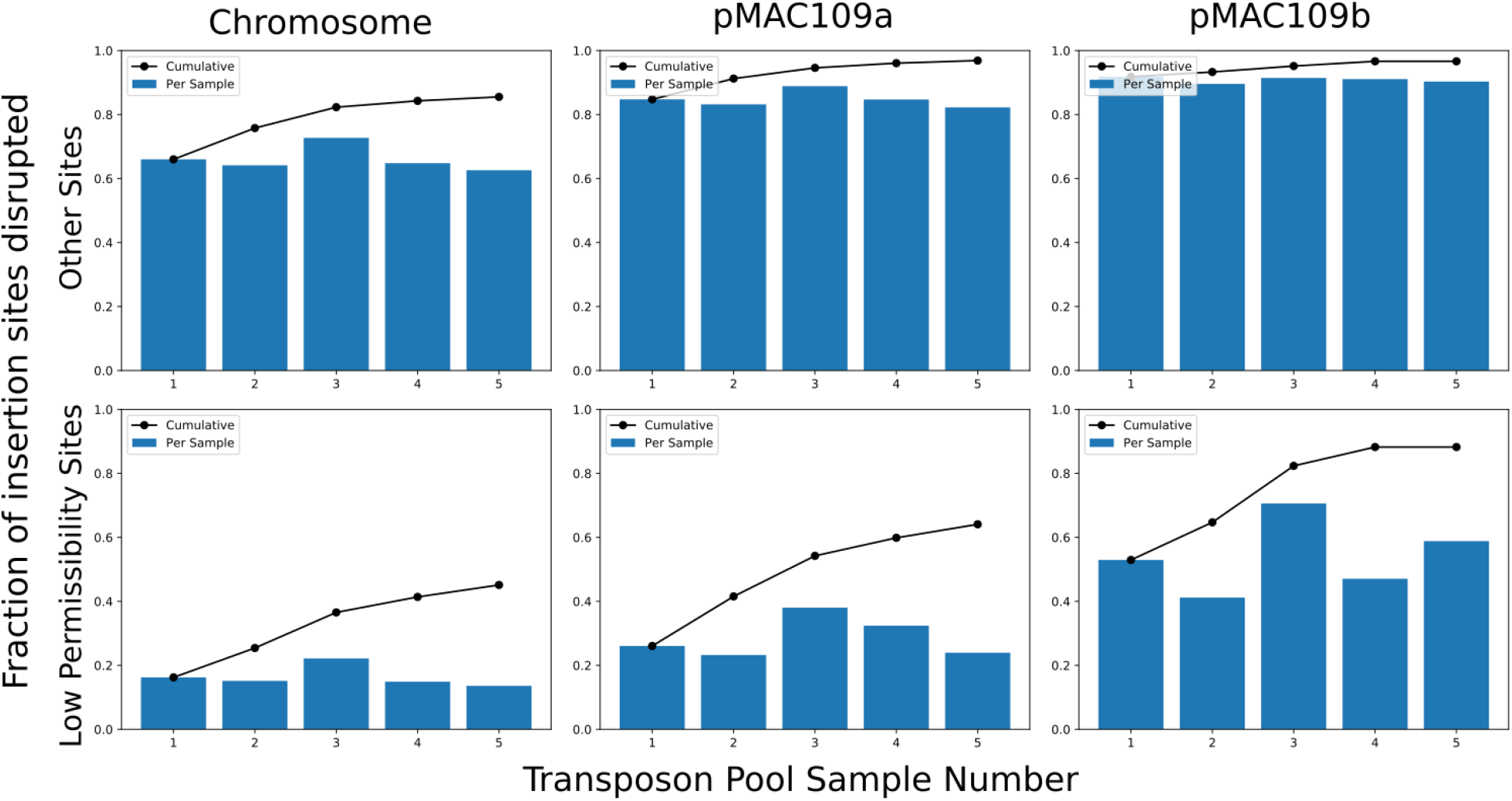
Each barplot shows the fraction of potential Himar1 insertion sites (TA dinucleotide) observed to have sustained at least one insertion in each independent pool of mutants for each replicon of the MAC109 genome. The line plots indicate the cumulative fraction of occupied insertion sites. Notably, the fraction of unique sites occupied saturates for sites not matching the previously defined sequence motif for low permissibility sites ([CG]GNTANC[CG]). However, sites matching this motif can be seen to be near saturation only in the case of the small plasmid (pMAC109b).

### Annotation of MAC109 Genetic Features

Our analysis method classified 270 features as ES, 489 features as GA, 1267 features as GD out of 5091 total annotated features. 73 features contained no TA sites and 9 features only contained TA sites shared with another feature. Therefore, these 82 features could not be evaluated with the Himar1 system. Our method classified 259 annotated coding sequences, 8 tRNAs, and 2 rRNAs as well as the only annotated tmRNA as essential. No annotated pseudogenes were labelled as essential though a minority of them were found to affect growth (i.e. were GA/GD). A summary of classifications by feature type is provided in Table 1 with classifications for individual features provided in Table S2. Table S3 provides these classifications merged with the raw read count data. Interestingly, our method identified 3 annotated coding sequences in pMAC109a and 1 coding sequence in pMAC109b as essential.

**Table 1:** Table of features annotated by our analysis method. NE = No Effect, GD = Growth Defect, ES = Essential, GA = Growth Advantage, N/A = Feature lacks potential insertion sites (TA dinucleotide) for the Himar1 transposon or only contains sites shared with another feature.

|  | CDS | Pseudogene | tRNA | Riboswitch | rRNA | ncRNA | tmRNA |
| --- | --- | --- | --- | --- | --- | --- | --- |
| NE | 2850 | 117 | 8 | 5 | 2 | 1 | 0 |
| ES | 259 | 0 | 8 | 0 | 2 | 0 | 1 |
| GD | 1208 | 32 | 26 | 0 | 0 | 1 | 0 |
| GA | 460 | 29 | 0 | 0 | 0 | 0 | 0 |
| N/A | 64 | 13 | 4 | 1 | 0 | 0 | 0 |

### Comparison of annotations with previously published transposon-based annotations

We compared the results of our analysis method applied to a previously published Tn-Seq dataset using *Mycobacterium tuberculosis* strain H37Rv^7^. All genes labelled as “ESD” (containing an essential domain) in the previously published dataset were considered essential for comparison. Figure 2 shows the overlap in the predicted essential coding sequences (CDS) from each method (RNA and other features excluded). Overall, there was good agreement between each method though our method appears to be somewhat more sensitive for essential gene detection than the previous method at this sample size. Upon inspection it was observed that the essential genes unique to our method contained a significant number of sites with zero or very few insertions, but these sites were interspersed among sites containing larger numbers of reads. This fits with expectations that the hidden Markov model used previously is sensitive primarily to multiple adjacent sites with low read counts, whereas our method is sensitive to the number of sites per gene regardless of adjacency.

**Figure 2:**
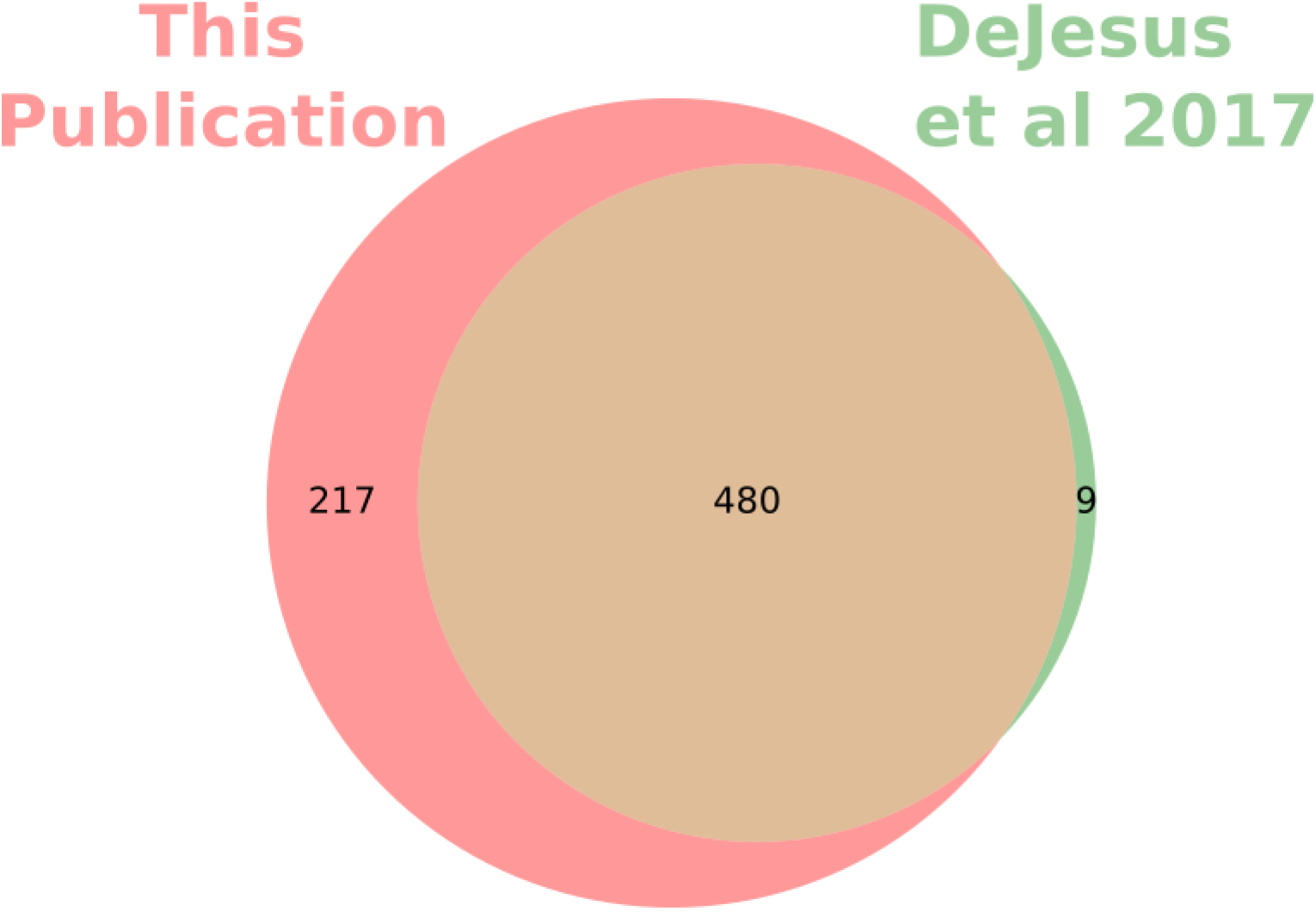
Venn diagram of essential genes predictions for *Mycobacterium tuberculosis* strain H37Rv from our analysis compared to the previously published essential gene predictions from DeJesus et al^7^. Notably, the genes labelled essential by the HMM are nearly a subset of the genes labelled as essential by our method. Only protein coding sequences are considered in this diagram.

## Discussion

We identified 230 genes as essential in both Mav and Mtb (Table S5). These may represent particularly good targets for drug development, as inhibitors of a gene product are likely to be effective against a close ortholog. As expected, a number of well-demonstrated targets are present. This includes the targets of the mycobacterial drugs cycloserine (alanine racemase, D-alanine – D-alanine ligase), rifamycins (RNA polymerase beta subunit), macrolides (50S ribosome), aminoglycosides (30S ribosome), fluoroquinolones (type IV topoisomerases and gyrases), bedaquiline (ATP synthase), and ethambutol (arabinosyltransferase). Additional compounds that have been reported to have some activity against mycobacteria include tryptophan synthase inhibitors^11^, ClpP inhibitors^12^, and Rho inhibitors (albeit only shown to be effective through genetic manipulation)^13^. A brief literature search also reveals many compounds that inhibit non-mycobacterial orthologs of these genes but appear to lack published results for killing activity in mycobacteria including inhibitors of GroEL^14^, RibBA^15^, SecA^16^, and LigA^17,18^. It is thus apparent that many opportunities are available for targeting these overlapping essential mycobacterial genes.

Our analysis classified four protein-coding genes on the two plasmids as essential (3 on pMAC109a and 1 on pMAC109b). This was somewhat surprising, as these plasmids have multiple copies per cell, and a disruption of a single gene copy should, in theory, be complemented by other copies. We used NCBI BLAST to find homologs of these genes. DFS55_24645 (on pMAC109a) and DFS55_25425 (on pMAC109b) are homologous to Rep, a protein critical for the replications of plasmids. Thus, one possible explanation for the essentiality of these Rep homologs is that plasmid copy number will decrease in daughter cells inheriting the plasmid (with no plasmid replication possible in a cell with all copies containing disrupted Rep). This is a strong selective pressure against the mutant plasmid. DFS55_14680 (on pMAC109a) is a ParA homolog. ParA controls the distribution of plasmids to daughter cells such that cells inherit the plasmid more equally. It is not immediately apparent how a more random distribution of the plasmids due to disruption of ParA would lead to a growth defect. Lastly, DFS55_24600 (on pMAC109a) is a hypothetical protein also classified as essential. It lacks a paralog in the MAC109 chromosome and an ortholog in Mycobacterium avium strain 104 (which does not contain plasmids). Thus, it appears to be non-essential for the *Mycobacterium avium* subsp. *hominissuis* pangenome. DFS55_24600 is homologous to Rv3081 from H37Rv and our analysis identified Rv3081 as “GD” (approximately 0.25 Relative Fitness). It is also apparent from examining the raw Tn-seq read counts (Table S3) that transposon insertion in the beginning of this gene does not have a profound effect on growth rate in MAC109 (this trend is less clear in H37Rv). Given these observations we can only speculate that this gene is addictive in MAC109 (and weakly addictive in H37Rv) – and may represent a toxin-antitoxin fusion with the toxin domain near the N-terminus. Future work could clone DFS55_24600 into an episomal (non-integrating) mycobacterial shuttle vector (such as pPB10) and examine the retention of the episome with and without this gene in the absence of antibiotic selection. Additionally, an attempt could be made to isolate a MAC109 mutant cured of pMAC109a.

Our analysis method has several advantages over other methods, including its anticipated high robustness as the result of using the zero-inflated negative-binomial to model read counts, which can more accurately account for non-saturating libraries, as these have a high probability of a site having no observed insertions. This may be especially important for transposons which cannot easily achieve saturation without very large numbers of transformants (due to lack of the strict TA site bias of Himar1), such as the Tn5 system^19^. Also, we have fully exploited the statistical independence of samples, which increases our statistical power. Other models, such as hidden Markov models, generally pool samples, limiting the usefulness of having biological replicates. However, our method also has limitations. Using our collected data, we detected a somewhat low number of essential features in MAC109 relative to H37Rv (270 and 738, respectively) despite evidence that the genome was saturated with insertions (Figure 1). Most likely, this is due to our somewhat low sample size (5 independent libraries). Therefore, we believe that sequencing additional independent transposon mutant libraries could significantly increase the statistical power to detect essential genes in MAC109, particularly for features with fewer insertion sites. A previous study^7^ used 14 independent libraries for H37Rv, which seemed to give our method good statistical power and may be a useful sample-size target for future studies. Additionally, while our method can correctly handle sites with low rates of insertion (e.g., [CG]GNTANC[CG]) it is possible that additional such sites exist that have not yet been defined. Defining the sites with low rates of insertion is especially important to avoid features falsely classified as essential.

In conclusion, we have generated genome-wide transposon mutant pools in *Mycobacterium avium* strain MAC109, collected sequencing data, and used a novel approach for annotating genes based on this data. We find that these pools are nearly saturated with transposon insertions, although not at low permissibility sites previously shown to have a reduced insertion rate. Our analysis identified the essential genes of MAC109 and we suggested explanations for the apparent detection of essential genes in the plasmids. We recommend that additional independent MAC109 transposon mutant libraries be collected, which we expect will greatly increase sensitivity. Future work could confirm our growth predictions by adapting the existing mycobacterial dCas9 knockdown system^20^ to *Mycobacterium avium* and measuring the impact of gene knockdown on bacterial growth rate.

## Materials and Methods

### Strains

MAC109, MAC104, OSU3388 were a gift from Dr. Luiz Bermudez (Oregon State University). MAC101 (Chester, ATCC 700898) was a gift from Dr. Eric Nuermberger (Johns Hopkins School of Medicine). Individual colonies of each strain were isolated and regrown to make stocks used in the described experiments. MAC101 was seen to form both translucent and opaque colonies. Both an opaque (MAC101o) and a translucent (MAC101t) colony were isolated and used for stocks.

ΦmycomarT7 was propagated and titered as previously described^21^. Final titers used for transformations exceeded 10^11^ PFUs/mL.

### Media and Buffers

To make 7H11 agar 10.25 grams of 7H11 w/o Malachite Green powder (HiMedia Cat No. 511A) was added to 450mL deionized water. 5mL 50% glycerol was then added before autoclaving. Hot agar was cooled to 55°C before addition of 50mL OADC enrichment and 1.25mL 20% Tween-80.

To make 7H9/10% OADC: 2.35g 7H9 powder was added to 450mL deionized water. After sterilization (via autoclaving at 121°C or by passing through a 0.22um filter) 50mL of OADC enrichment (Becton-Dickinson) was added. Unless otherwise specified, no Tween-80 or glycerol was included.

To make 7H9/50% OADC: Recipe identical to 7H9/10% OADC but using 250mL water and 250mL OADC.

To make PBS-Tw: 1.25mL filter-sterilized 20% Tween-80 was added to 500mL sterile PBS.

To make MP Buffer: 50mM Tris-HCl (pH 7.5), 150mM NaCl, 10mM MgSO4, 2mM CaCl2. Autoclave individual components before combining.

### Testing of transformation efficiency of Mav strains

Five strains of Mav (MAC109, MAC104, OSU3388, MAC101o, MAC101t) were tested for transformation by ΦmycomarT7. For transformation, strains were grown in 150mL of 7H9/10% OADC. After OD of each strain reached 0.32 – 0.89, 100mL of cultures were equally split into two 50mL conical tubes. Bacteria were pelleted via centrifugation and resuspended in 10mL MP buffer. Bacteria were pelleted again via centrifugation and resuspended in 4.5mL MP Buffer. 0.5mL of MP Buffer (negative control) or ΦmycomarT7 stock (approximately 10:1, phage:bacteria) was added to each tube. Tubes were incubated for two days shaking at 37°C. Bacteria were then pelleted via centrifugation and resuspended in PBS-Tw (phosphate-buffered saline containing 0.05% Tween-80). Bacteria were then spun down again and resuspended in 1mL of PBS-Tw. Transformed bacteria and negative control for each strain were then diluted in PBS-Tw and plated on 7H11 with and without 50ug/mL kanamycin for titration.

### Generation of transposon mutant libraries in MAC109

In preliminary experiments, we found that MAC109 growth increased at higher concentrations of OADC. We suspect the oleic acid in OADC is the key to achieving this, based on previous reports^22^. 5 independent transposon mutant pools were generated. MAC109 was grown in 700mL 7H9/50%OADC to OD 2.1 in two 1.5L roller bottles shaking at 37°C. Based on previous results (data not shown) we estimated the initial bacterial density based on optical density to be 4 × 10^8^ CFUs/mL for calculation of volume of phage stocks. Bacteria were aliquoted to 12-50mL conical tubes and centrifuged (2000g for 5 minutes) and supernatant removed. 5mL MP Buffer was added to each tube and bacterial pellet was resuspended. Pairs of tubes were pooled yielding 6-10mL aliquots. Samples were then centrifuged (2000g for 5 minutes) and supernatant removed. Phage (10:1, phage:bacteria) was then added to all tubes except no-vector control. MP Buffer was added to all tubes to a final volume of 5mL and bacterial pellets were dispersed via pipette. Bacterial/phage mixtures were then placed on a shaker incubator (37°C) for two days. Tubes were then centrifuged (2000g for 5 minutes) and supernatant removed. 10mL PBS-Tw was then added and the bacterial pellet was dispersed via pipette. Tubes were then spun down again (2000g for 5 minutes), supernatant removed, and 1 mL of PBS-Tw was used to resuspend pellets.

50uL of each tube of washed transformants (or no-vector control) were diluted and plated on 7H11 plates, with or without 50ug/mL kanamycin, to determine transformation efficiency and background resistance. The remainder of the cultures were plated on 7H11 containing 50ug/mL kanamycin in Pyrex baking dishes (15” x 10”, 500mL agar per dish, 1 tube per dish). After 7-10 days colonies were scraped from each dish and dispersed in fresh 7H9 broth and frozen in aliquots at -80°C for later use.

DNA was extracted from one aliquot of each transposon mutant pool using a previously described gDNA extraction protocol for short read sequencing^10^. We adapted a previously published library prep protocol^23^ to prepare libraries for sequencing. Adaptations include the use of magnetic beads for purification and library size selection as well as changes to PCR conditions (for details see Text S1). Libraries were sequenced (2×75bp) on an Illumina HiSeq 2500 by the Johns Hopkins GRCF High Throughput Sequencing Center. 5 independent libraries were sequenced yielding between 2,194,085 – 4,381,545 reads per library for a total of 18,197,728 paired-end reads.

### Raw Data processing

We previously showed that the MAC109 genome contains two plasmids in addition to the bacterial chromosome. We adapted the TRANSIT pre-processor (tpp)^24^ to allow for mapping to multiple contigs. These changes were included in the release of TRANSIT/tpp v2.4.1. We used tpp v2.4.1 to map all reads to the MAC109 genome. Command for processing raw reads: tpp -himar1 -bwa -bwa-alg aln - ref MAC109.gb -replicon-ids “CP029332,CP029333,CP029334” -reads1 TnPool_1.fastq -reads2 TnPool_2.fastq -window-size 6 -primer AACCTGTTA -mismatches 2. After PCR duplicate removal, a total of 10,597,261 unique reads mapped to the genome and were used for analysis.

### Statistical Analysis

We use a previously suggested labelling scheme^25^ to annotate each gene of MAC109. A gene is labelled NE (No Effect) if a transposon insertion in any of its potential insertion sites causes no effect on growth. A gene is labelled GD (Growth Defect) if it contains at least one insertion site such that upon transposon insertion it results in a decrease in bacterial growth. A gene is labelled GA (Growth Advantage) if it contains at least one insertion site such that upon transposon insertion it results in an increase in bacterial growth. A gene is labelled ES (essential) if it contains at least one insertion site such that upon transposon insertion it results in a large loss in viability.

To annotate the MAC109 genome, we have designed a robust procedure. Some additional details of this method are provided in the supplement (Text S2). At a conceptual level, our analysis pipeline proceeds in two steps. First, insertion sites without a growth defect are approximately identified with a rank-based filter procedure. Second, the counts at the insertion sites identified by the filter are assumed to approximate the null distribution and used for statistical hypothesis testing. For identification of ES genes, the approximate null distribution is fit to a zero-inflated negative binomial distribution (using maximum likelihood estimation) which is then scaled and used for hypothesis testing. For identifying the GD and GA sites, the empirical cumulative distribution function is used for hypothesis testing. Stouffer’s method is used to combine p-values from multiple replicates and multiple sites. Lastly, multiple hypothesis correction is performed (Benjamini-Hochberg for ES, Bonferroni for GD/GA testing).

#### Relative Fitness

The fitness, relative to wildtype, resulting from disruption of a particular gene is approximated as follows. First, the mean of the read counts at each insertion site is calculated across samples. The site fitness is calculated as the mean read count of each site divided by the median across all sites (i.e., samples are normalized to the median). Finally, each gene is assigned a Relative Fitness equal to the median of the site fitness for all sites contained in the gene.

#### Rank-based filter procedure

We assumed that all mutants with a transposon insertion at the same site will have identical growth rates (i.e., the growth rate is entirely defined by the insertion site). We also assumed that not more than 40% of insertion mutants would have a growth defect and not more than 15% of mutants would have a growth advantage (and therefore at least 45% of mutants would have a growth rate that is identical to wildtype). We selected these thresholds based on previous predictions in *Mycobacterium tuberculosis*^7^ suggesting that 15% of insertion sites cause a growth defect and 8% cause a growth advantage. We have added a large margin of error to ensure conservatism.

Note that if some of the identities of insertions mutants with growth rates identical to wildtype were known ahead of time we could simply use the distribution of the reads at these sites to train a null model to test the other sites. This is the intuition behind our rank-based filter procedure. However, as the identities of the insertion sites with no effect on growth rate are unknown we use an approximation. For each of *J* transposon pools (replicates) we compute the rank of the read count at each site (averaging identical ranks) in the other *J*-1 samples. For each site, we then take the average of these J-1 ranks across samples. Lastly, we order the average rank from least to greatest and remove the smallest 40% and greatest 15% (removing additional sites with ties at the threshold), leaving only ~45% of the original insertion sites. The read counts from these remaining ~45% of sites will be distributed approximately the same as an insertion site with no effect on growth. Additionally, previous literature suggests that the Himar1 transposon is biased against insertion sites with the motif (GC)GNTANC(GC)^7^. Therefore, we separately apply the above rank-based filter to the read count data collected from these sites.

To demonstrate the correctness of our rank-based filter procedure we utilized simulated data. Briefly, read counts from 39,000 insertion mutants without a defect were simulated as a negative binomial distribution with mean 35 and dispersion 3.0. These parameters are roughly those found by fitting real data (fitting procedure described below). Additionally, read counts from 15,000 mutants with a growth defect were simulated with a mean of between 0 and 0.67 times that of a no defect mutant using the negative binomial distribution with an identical dispersion. The mean multiplier was chosen for these mutants by uniform sampling between these bounds. Lastly, read counts from 6,000 mutants with a growth advantage were simulated using 1.5 to 4 times the null mean (uniformly distributed) and identical dispersion. Combining these 3 groups of samples provided a simulated transposon mutant library. 5 and 50 independent simulated transposon mutant libraries were generated. The rank-based filter procedure described above was then applied to the resulting datasets. Q-q plots provided in Figures S1B and S1D comparing the theoretical distribution to the unfiltered and filtered empirical cdfs show that the filter procedure improves accuracy. Increased sample size also improves accuracy, as expected.

#### Hypothesis Testing for Essentiality (ES)

To classify a gene as ES, we performed statistical hypothesis testing. The read counts from the insertion sites identified by the rank-based filter are used to fit a zero-inflated negative binomial distribution (See Text S2 for definition). Fitting is done by maximizing the likelihood with L-BFGS-B as implemented in scipy.optimize (Scipy v1.2.1). Using the fit distribution, we then create a new “borderline ES” distribution by scaling the mean of the negative binomial distribution to 5% of the original, keeping the dispersion and zero inflation component constant. We use this borderline distribution to do statistical hypothesis testing on the read counts from each of the sites using the lower tail probability as the p-value. This means that a gene whose insertion gives 5% of WT growth is unlikely to be called ES. While the particular threshold we have chosen (5% of wildtype growth) is somewhat arbitrary, we feel it is both small enough to ensure mutants labelled ES are highly defective but not so small so as to have no hope of classifying highly defective mutants as ES.

To pool essential p-values across samples, we used the one-tailed Stouffer’s method at each site. To pool p-values across insertion sites within a gene we use the truncated product method^26^ with a truncation threshold of 0.5 (τ < 0.5). TPM provides a principled approach for limiting the effect of sites with no associated growth defect which would otherwise greatly inflate the p-values (such as those sites at the C-terminus of the gene which may not disrupt the function of the protein). We then control the False Discovery Rate (FDR) using the Benjamini-Hochberg procedure (FDR < 0.01).

#### GD/GA Hypothesis Testing

To classify a gene as GD or GA, we performed statistical hypothesis testing. We utilized the read counts for insertion sites identified by the rank-based filter to form an approximate null distribution and used the empirical cumulative distribution function (ecdf) to compute p-values. We generated a separate ecdf for low permissibility sites. We also generated separate ecdfs for each contig as sequencing depth varied greatly between contigs (due to multiple copy-number plasmids). The exact p-value computation, which ensures p-values are continuously distributed, is described in detail in the supplement. For a particular insertion site, the p-values from each sample were pooled using the one-tailed Stouffer’s method. The resulting pooled p-values from all insertion sites within the same gene were then pooled using the two-tailed Stouffer’s method. For declaring genes as GA or GD we set the p-value threshold to allow only a single (expected) false discovery after 5009 tests, corresponding to a single-test p-value of approximately 0.0002. A gene was declared GD if its Relative Fitness was less than 2/3 and was statistically significant (p < 0.0002). Similarly, a gene was declared GA if its Relative Fitness was greater than 1.5 and was statistically significant at the same threshold. Note that if a gene meets the criteria for both the GD and ES label then it is given the ES label only. If it meets the ES criteria but not the GD label it is given the NE label.

### Data and Source Code Availability

We have made efforts to enable others to reproduce the major results of this paper from the raw data. Scripts and instructions for use are provided at GitHub (https://github.com/joelbader/essential_genes)^27^. Raw data is provided in NCBI’s SRA under accession number: PRJNA527645.

## Acknowledgements

This publication was made possible by support from the Sherrilyn and Ken Fisher Center for Environmental Infectious Diseases, Division of Infectious Diseases of the Johns Hopkins University School of Medicine. Its contents are solely the responsibility of the authors and do not necessarily represent the official view of the Fisher Center or Johns Hopkins University School of Medicine.

We are grateful to Dr. Luiz Bermudez for providing strains and advice during this project.

---

**Figure S1:** Simulated data showing correctness of rank-based filter. As described, simulated read counts were generated to test our rank-based filter procedure. A simulation of either 5 sequenced samples ((A) and (B)) or 50 samples ((C) and (D)) was generated. (A) and (C) show histograms of the read counts across all sites before applying the filter procedure in green and the read counts after applying the filter procedure in blue. In red we plot the pmf of our sampling distribution for the null distributed sites. Performance was assessed by q-q plots in (B) and (D). In green are the empirical quantiles before applying the rank-based filter procedure and in blue are the quantiles after filtering. The red line represents perfect theoretical performance.

## Descriptions of supplemental files

- Supplemental Figure S1: Simulated data showing correctness of rank-based filter
- Supplemental Text S1: Protocol for preparing sequencing libraries
- Supplemental Text S2: Additional details of analysis method
- Supplemental Table S1: Transformation efficiency of avium strains.
- Supplemental Table S2: Gene prediction in MAC109 (with p-values and LFC)
- Supplemental Table S3: Raw data in MAC109 along with gene predictions
- Supplemental Table S4: Essential genes in H37Rv based on previous data
- Supplemental Table S5: Overlap between MAC109 and Mtb essential genes (both computed with our analysis method)

